# On how the dentate gyrus contributes to memory discrimination

**DOI:** 10.1101/235473

**Authors:** Milenna T. van Dijk, Andre A. Fenton

## Abstract

**Summary:** The dentate gyrus (DG) is crucial for behaviorally discriminating similar spatial memories, predicting that dentate gyrus place cells change (“remap”) spatial tuning (“place fields”) for memory discrimination. This prediction was never tested, although DG place cells remap across similar environments without memory tasks. We confirm this prior finding, then demonstrate that DG place fields do not remap across spatial tasks that require DG-dependent memory discrimination. Instead of remapping, place-discriminating discharge is observed transiently amongst DG place cells, particularly where memory discrimination is most necessary. The DG network signals memory discrimination by expressing distinctive sub-second network patterns of co-firing amongst principal cells at memory discrimination sites. This is accompanied by increased coupling of discharge from excitatory principal cells and inhibitory interneurons. Instead of remapping, these findings identify that memory discrimination is signaled by sub-second patterns of correlated discharge within the dentate network.

**eTOC blurb:** van Dijk and Fenton report that dentate gyrus place cells signal memory discrimination not by remapping, but by variable sub-second patterns of coordinated place cell network discharge and enhanced discharge coupling between excitatory and inhibitory neurons, at sites of memory discrimination.

**Highlights:**

- Dentate gyrus-dependent memory discrimination does not require place cell remapping
- Dentate neural correlates of pattern discrimination are transient, lasting seconds
- Sub-second dentate network discharge correlations signal memory discrimination
- Dentate excitatory-inhibitory coupling is increased at memory discrimination sites

## Introduction

Hippocampus is crucial for discriminating between memories (Gilbert et al., 1998; Yassa and Stark, 2011), for navigating space (Maguire et al., 1998; Morris et al., 1982; O’Keefe and Nadel, 1978), and the discharge of hippocampus place cells represents locations (O’Keefe, 1976; Wilson and McNaughton, 1993). Although place cell discharge is unreliable in time (Fenton and Muller, 1998), the discharge reliably localizes to specific regions of space called the cell’s place field (O’Keefe, 1976). These place fields relocate when the environment changes sufficiently, a process known as “spatial remapping” (Muller and Kubie, 1987) or alternatively, they maintain their locations but systematically change firing rates in what is called “rate remapping” (Hayman et al., 2003; Leutgeb et al., 2005b). Both forms of remapping have been associated with memory discrimination (Alme et al., 2014; Colgin et al., 2008; Wills et al., 2005), and the dominant cognitive map theory predicts place fields will remap across conditions requiring distinct place memories (O’Keefe and Nadel, 1978).

Theories of hippocampus memory computation assert the DG is specialized for discriminative functions such that DG outputs are more distinctive than the corresponding inputs, what is called “pattern separation” (Marr, 1971; O'Reilly and McClelland, 1994; Treves and Rolls, 1994). As predicted, compromising DG function impairs difficult memory discriminations in modestly distinct environments (Burghardt et al., 2012; Gilbert et al., 2001; Kheirbek et al., 2013; Lee et al., 2005; McHugh et al., 2007; Nakashiba et al., 2012). In addition, DG principal cells (pDGC) consisting of granule cells and mossy cells remap in response to sufficiently large environmental changes (Danielson et al., 2017; GoodSmith et al., 2017; Leutgeb et al., 2007; Neunuebel and Knierim, 2014; Senzai and Buzsaki, 2017), but the extent to which these observations indicate pattern separation and whether they are relevant to memory discrimination is not established.

Indeed, pDGCs place cells have been characterized but not during an explicit memory discrimination task that requires intact DG function (Danielson et al., 2017; GoodSmith et al., 2017; Leutgeb et al., 2007; McHugh et al., 2007; Neunuebel and Knierim, 2014; Senzai and Buzsaki, 2017). To critically test the prediction that DG remapping mediates memory discrimination we recorded DG neurons while mice on a rotating arena perform DG-dependent memory discriminations in active place avoidance tasks (Burghardt et al., 2012; Kheirbek et al., 2013; Park et al., 2015). The rotation dissociates the environment into two spatial reference frames and reveals that distinct frame-specific patterns of ensemble place representations alternate in the ensemble spike time series (Fenton et al., 1998; Kelemen and Fenton, 2010). We find that DG place cells do not spatially remap across sessions that require DG-dependent memory discrimination, although rate remapping increases on memory trials. When the mice are in the vicinity of the avoided place, DG place cells preferentially discharge in the task-relevant spatial frame, indicating transient and distinctive network states. Instead of remapping, the precise locations where memory discrimination is required were signaled by sub-second co-firing of pDGC place cells that was inconsistent with the overall discharge correlations of the cell pair. This network inconsistency was accompanied by increased sub-second discharge correlations amongst DG place cell-interneuron pairs. Instead of the time-averaged changes in spatial tuning associated with remapping, these findings point to dynamic patterns of correlated DG discharge for memory discrimination.

## Results

### Confirmation that DG place fields are less stable than CA fields across environments

We began by recording cells across three versions of an environment (standard, 90° cue relocation, and wall removal; Fig. 1A, Fig. S1) to confirm that place cells in DG (45 place cells of 133 cells in 5 mice) are more sensitive to environmental changes than those in Ammon’s horn (20 place cells of 42 cells in 3 mice) (Leutgeb et al., 2007). The majority of pDGC (64%) and CA (75%) place cells had a single place field during the initial recording in the standard environment (test of proportions z = 1.6, p = 0.1). The number of place fields scaled with the 2-fold increase in area from the standard to the wall removal condition (DG standard: 1.42±0.10 fields; removed: 2.70±0.29 fields; CA standard 1.26±0.13 fields; removed 2.0+0.28 fields; region: F_1,63.7_ = 3.65; p = 0.10; condition: F_1,63.3_=13.4 p=10^−4^, interaction: F_1,63.3_ = 1.26 p = 0.27 post-hoc: removed DG > standard DG and CA; removed CA > standard CA), as previously reported for rat place cells (Fenton et al., 2008; Park et al., 2011). Other discharge properties were also similar for the DG and CA principal cell populations (Table S1). In contrast, pDGC firing rate map stability was less than in CA and the stability of both populations was changed by wall removal but not by the cue relocation (Fig. 1B). Rate remapping in pDGCs and CA cells did not differ across the environment manipulations (region: F_1,172_=0.16, p=0.68; manipulation: F_2,172_ = 2.8, p = 0.064; region x manipulation: F_2,172_ = 0.2, p= 0.81), but the magnitude of the firing rate changes in and out of the primary place field differed across manipulations for CA rates but not for DG rates (Fig. 1C).

**Figure 1.**
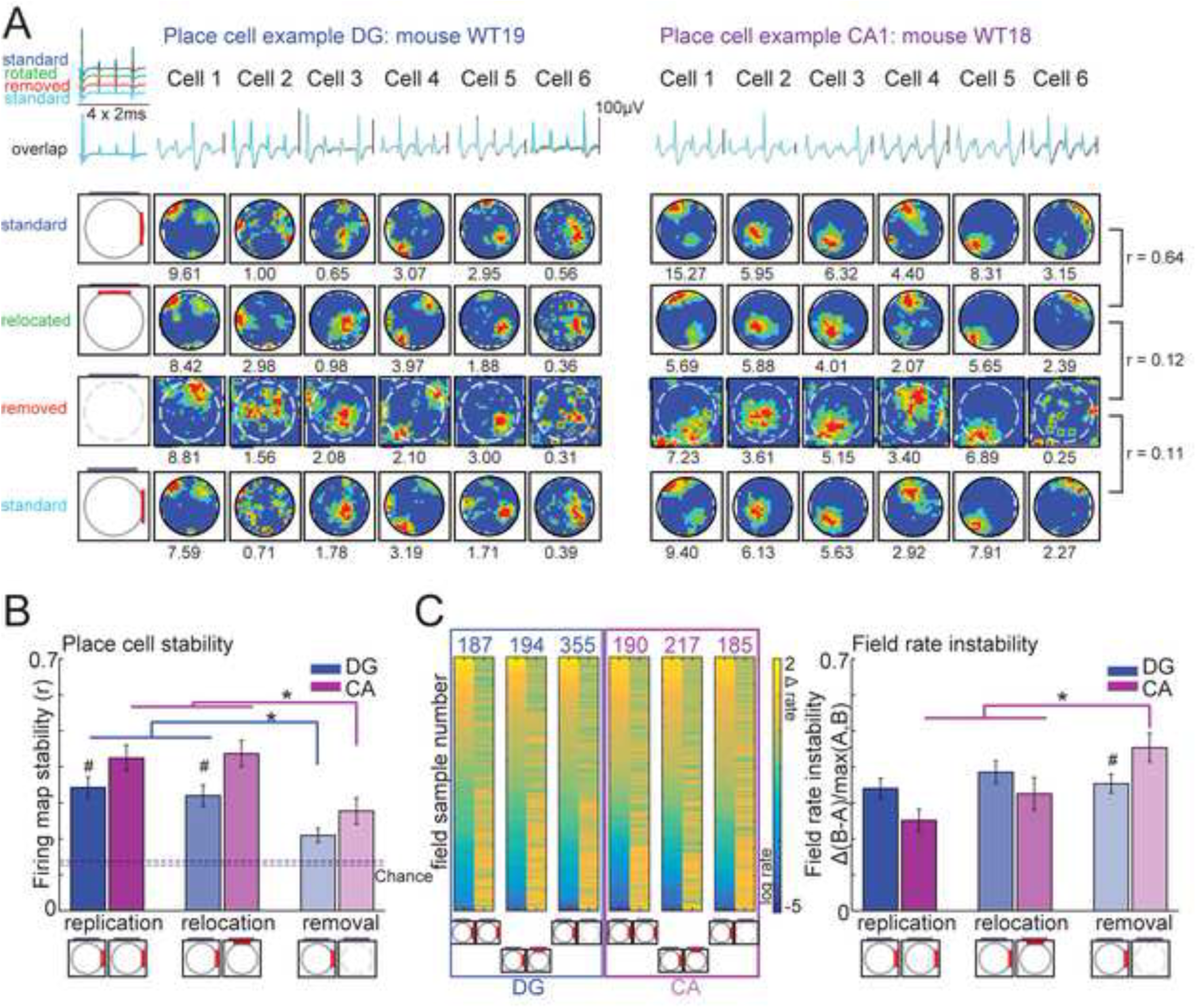
DG place cells are less stable than those in CA1/3 and place cells in both regions respond to changes to the environment. a) Example DG (left) and CA1 (right) firing rate maps of mice exploring three versions of an environment: a transparent cylindrical enclosure with cue cards in a square box with a cue card (*standard*), the same cylindrical enclosure rotated 90° while the box remained stable (*rotated*) and the square box by itself (cylinder *removed*). b) Firing map stability measured by correlating rate maps of each cell across a pair of conditions. We define three comparison manipulations: *replication* = [standard-standard (illustrated), rotated-rotated, removed-removed], *rotation* = [standard-rotated] and *removal* = [standard-removed (illustrated), rotated-removed]. Stability of place cells in Ammon’s horn (CA) is higher than those in DG. Stability is lower for the removal condition than for replication and rotation conditions (region: F_1,172_ =15.1, p = 0.0001; condition effect F_2,172_=12.4, p<0.0001; interaction: F_2,172_, p=0.77). c) Left: Field rates and rate changes for the first versus second trial of each manipulation. The magnitude of field rate changes differed between manipulations in CA where the removal manipulation showed larger changes than replication and rotation and than removal in DG (region: F_1,334_ = 0.16, p = 0.69, manipulation: F_2,334_ = 5.43, p = 0.0048, interaction: F_2,334_ = 3.91, p = 0.021). These results confirm that DG place cells are sensitive to changes in the environment and that DG shows less firing rate map stability than CA. See also Figure S1. Bar graphs represent mean ± SEM. * signifies regional differences, # signifies manipulation differences.

### Confirming a DG-dependent memory discrimination task

Before testing the prediction that place fields change with memory discrimination, we confirmed that the active place avoidance memory discrimination task depends on DG function (Fig. 2A). Laser illumination of POMC-Halorhodopsin mice that optogenetically silences granule cells in the Cre^+^ but not Cre^−^ littermates (n’s = 8) showed that DG cells are not essential for learning to avoid the initial location of shock, but are essential for avoiding shock on the conflict trial when the shock is relocated 180° (Fig. 2A,B). Because illumination both directly and indirectly silences DG cells (Senzai and Buzsaki, 2017) this demonstrates the DG is specifically necessary for conflict memory discrimination (Burghardt et al., 2012; Kheirbek et al., 2013).

**Figure 2.**
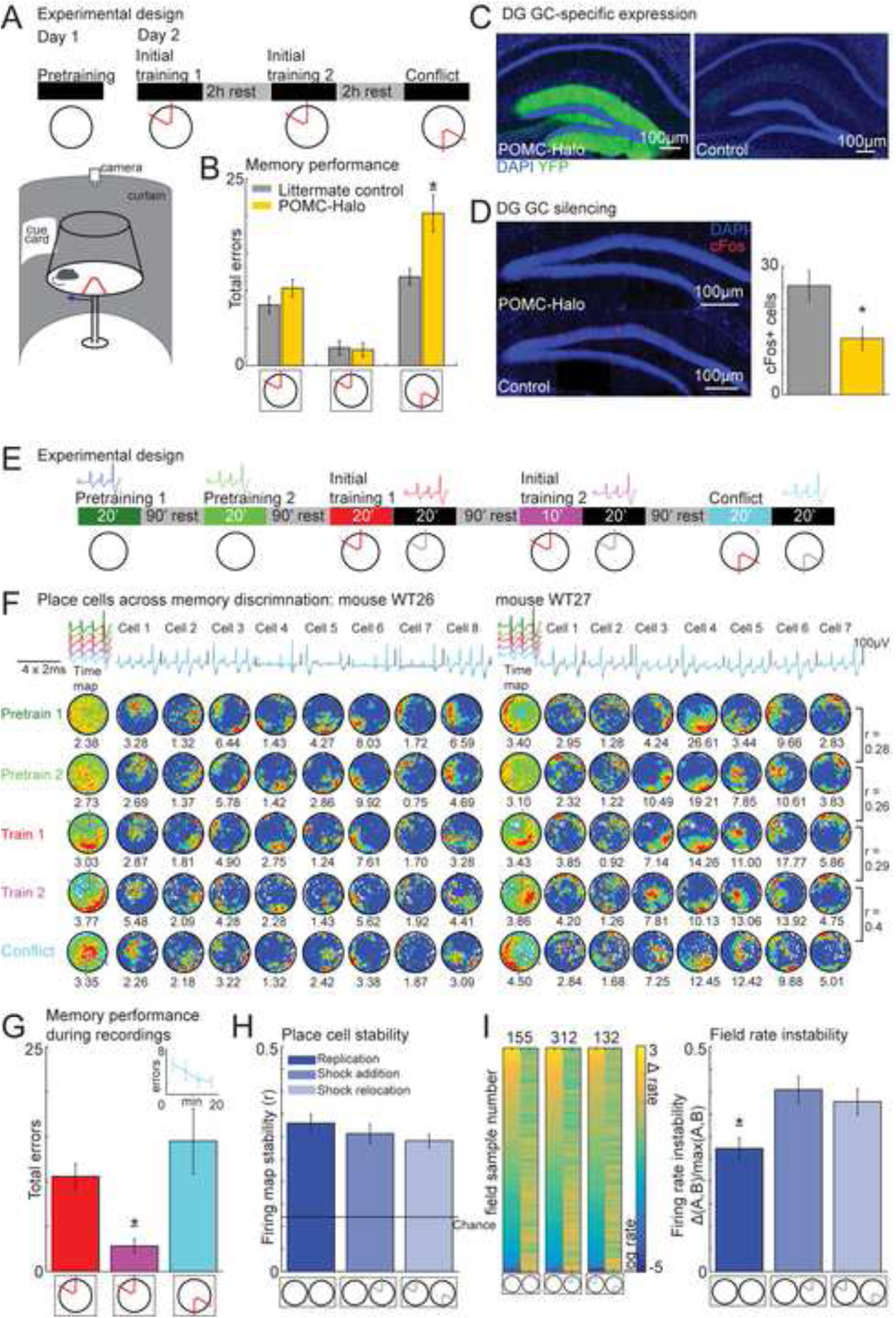
Memory discrimination occurs independently of spatial remapping but DG cells modulate their rates in response to task change. a) Behavioral protocol. Day 1: 30-minute pretraining trial (shock off, 66.7±10.7 entrances (“errors”)) on the rotating arena. Day 2: mice learn in two 30-minute initial training trials (2 hours apart) to avoid a shock zone that is stationary with respect to the room. During the conflict trial the shock zone is relocated 180° and mice learn to distinguish between the current and previous shock zone locations. b) Mice with light-silenced granule cells learn avoidance at similar rates as control mice, but after the shock zone relocation, Cre^+^ POMC-Halorhodopsin mice are deficient in discriminating between the old and new shock zone location (group: F_1,14_=11.7, p = 0.004; condition: F_1.42,19.9_=46.4, p<0.0001; interaction: F_1.42,19.9_ = 4.9, p = 0.03), confirming that intact DG function is necessary for memory discrimination in this paradigm. c) Expression of halorhodopsin is limited to DG GCs in Cre^+^ POMC-Halorhodopsin mice and is not expressed in Cre^−^ littermate controls. d) Cre^+^ POMC-Halorhodopsin mice show fewer c-Fos^+^ GCs labeled during the conflict trial (t = 2.7, p = 0.04). e) Recording protocol. Day 1: two pretraining sessions (shock off, 68.8±10.6 entrances) on the rotating arena, followed by two training sessions, then memory discrimination is tested in the conflict trial in which the shock zone is relocated 180°. Mice were trained and immediately after the training or conflict trial the shock was turned off and single units were recorded. Rest in the home cage between trials was 90 min. f) Example DG firing rate maps from two mice during each session. Each unit’s average concatenated tetrode waveform for each session is overlaid above each rate map. Dwell maps show the time spent in each pixel for each session. g) The mice learned to avoid in the first training trial and maintained their memory during the second trial. Errors in the conflict trial initially increased in response to the new shock location then quickly decreased as shown by the learning curve in the inset (condition: F_2,4_ = 18.2, p=0.01, training 2< training 1 and conflict; Errors: training 1: 10.6±1.4, training 2: 2.8±0.8, conflict: 14.5±3.6). h) Firing rate map stability did not differ between the replication (pretraining-pretraining [illustrated], training 1-training 2), shock addition (pretraining-training [illustrated]) and shock relocation (training-conflict [illustrated]) manipulations (H_2,117_ = 1.02, p = 0.60) and was no different from the replication condition in Fig 1B. i) Left: Field rates and rate changes for the first vs. second trial of each manipulation. Rate remapping is greater for the shock addition and shock relocation manipulations compared to the replication manipulation (H_2,237_ = 13.25, p<0.0013). Bar graphs represent mean ± SEM. * indicates different as shown by lines; * indicates significantly different from all other manipulations.

### DG-dependent memory discrimination is not accompanied by dentate firing field remapping

We then used a similar protocol (Fig. 2C) to test if pDGC firing rate maps (n = 42 place cells of 113 cells in 6 wild-type mice) change with memory discrimination (Fig. 2D). Place avoidance learning was normal (Fig. 2E) and measures of spatial firing quality did not differ across the pretraining, initial training, and conflict training trials (Table S2). Importantly, firing rate map stability also did not differ across the “replication manipulation” (pretraining 1 vs. pretraining 2, initial training 1 vs. initial training 2), across the “shock addition” manipulation (pretraining vs. initial training) and across the “shock relocation” manipulation (initial training vs. conflict training; Fig. 2F). These estimates of stability were similar to the estimates across the replication manipulation of the environment (Fig. 1B) for pDGC (z = 0.11, p > 0.9) but were weaker compared to CA cells (z = 2.6, p < 0.01). The magnitude of pDGC-specific firing rate changes (H_2,117_ = 14.4, p<0.001) and place field-specific rate changes were greater in response to the shock-added and shock-relocated manipulations than across the replication manipulations (Fig. 2G). This pattern of replication < shock addition = shock relocated was also observed in the putative inhibitory neurons (H_2,57_ = 8.97, p=0.011). These findings demonstrate DG firing rates are sensitive to changes in task contingency, consistent with episodic encoding (Leutgeb et al., 2005b). Note however that these single-cell estimates of neural pattern discrimination were not specific to DG-dependent memory discrimination.

### Dentate place cell ensembles transiently and purposefully discriminate places in distinct spatial frames

We next examined these DG data by analyzing the conjoint discharge of cells on subsecond time scales. We were motivated because the assumptions of the preceding session-averaged analyses of single-cell firing (Figs. 1,2) contrast with the strong non-stationarity (Carr and Frank, 2012; Fenton et al., 2010; Fenton and Muller, 1998; Ferbinteanu et al., 2011; Gothard et al., 2001 ; Gothard et al., 1996; Gupta et al., 2010; Huxter et al., 2003; Jackson and Redish, 2007; Redish et al., 2000; Shapiro and Ferbinteanu, 2006; Singer et al., 2010) and ensemble place coding properties (Dupret et al., 2010; Harris et al., 2003; O'Neill et al., 2008; Park et al., 2011; Pastalkova et al., 2008; Pfeiffer and Foster, 2015; Wikenheiser and Redish, 2015; Wilson and McNaughton, 1993) that are reported for hippocampal spiking dynamics. We thus considered a different form of neural pattern discrimination that can be measured on a moment-to-moment basis. The rotation of the arena dissociates the environment into two distinct spatial frames. One is stationary, defined by room-anchored cues and the other is rotating, defined by local arena-anchored cues (Fenton et al., 1998). To solve this task the mouse has to discriminate between room-based spatial information and arena-based spatial information and this is essential in the vicinity of the shock zone if the animal is to successfully avoid the shock that is defined only by room-frame information, and not by arena-frame information. Spatial frame-specific positional discharge was estimated by separately computing momentary positional information (*I_pos_*) in each spatial frame for the place cell ensemble (Kelemen and Fenton, 2010; Olypher et al., 2003), and computing *ΔI_pos_*, the difference between room-frame and arena-frame *I_pos_*each 133 ms (Fig. 3A). Like place cell firing rate fluctuations (Fenton et al., 2010), the correlation between *ΔI_pos_* and running speed explains little of the variance (r^2^=0.69%, p < 0.001). Room information dominates arena information more often than *vice versa* during pretraining but this preference disappears when shock is added or relocated (Fig 3B) even though room information must be used to avoid shock (Bures et al., 1997; Fenton and Bures, 2003). During the ~1 s before mice avoid or enter the initial location of shock (Fig. 3C) *ΔI_pos_*is positive (i.e. room-preferring) and larger than when the mice are in the corresponding area 180° away in the potentially safest zone (Fig. 3C). The prevalence of room-preferring information followed the shock zone relocation on the conflict trial (Fig. 3C), even though time-averaged firing rate maps did not change (Fig. 2). We observed a similar preference for room-specific discharge in the vicinity of the current shock zone when we examined the locations of frame-specific *I_pos_* values (Fig. 3D), instead of their differences. These findings demonstrate discrimination of neural representations of location in DG discharge; the discrimination is both dynamic and purposeful insofar that it corresponds to the need for discriminative place avoidance memory. Notably, place discrimination into room and arena-defined places was insensitive to optogenetic silencing of DG cells (Fig. 2B).

**Figure 3.**
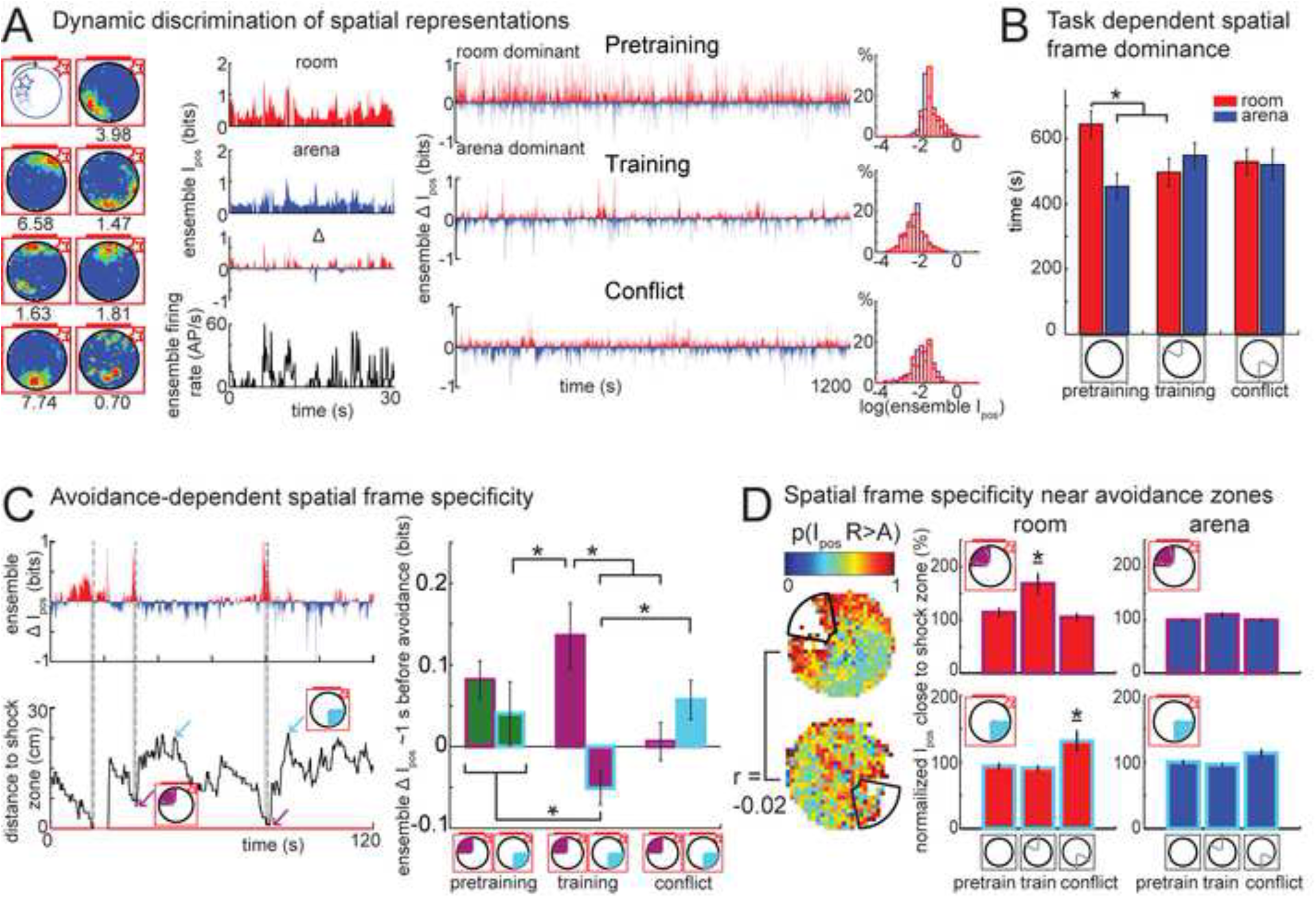
Multistable, purposeful DG place cell ensemble representations during place avoidance. a) Example place cells and corresponding room (top panel) and arena (second panel) ensemble *I_poS_* over time as well as *ΔI_poS_* (third panel) and ensemble firing rate (bottom panel). Right: Example of an ensemble’s *ΔI_pos_* time series during pretraining, training and conflict trials and the corresponding frame-specific *I_pos_* histograms. b) Room information dominates arena information more often in pretraining trials, but the room frame preference disappears when shock is added or relocated (spatial frame: F_1,10.6_,= 2.0, p = 0.18, condition: F_2,19.8_ = 0.3, p = 0.78; frame x condition: F_2,19.8_ =4.9, p = 0.018). c) Left: example of *ΔI_pos_* time series (top) during a 2-min period and the mouse’s concurrent avoidance behavior shown as distance from the shock zone (bottom). Right: After training or conflict trials, *ΔI_pos_* is positive (room-dominating) and larger during ~1s before active avoidance (defined as a local minimum in the distance to a shock zone time series) compared to when mice are in the safe zone that is the corresponding region on the opposite side of the arena or compared to when mice are close to that shock zone in trials that it was safe (condition: F_2,18.5_=0.8, p=0.47; shock zone: F_1,9.9_= 3.3, p = 0.10; condition x shock zone: F_2,18.5_=11.4, p=0.0006). d) Summary maps of the proportion of time mice were in the room-dominant state during training (top) and conflict (bottom) trials show a preference for preferentially signaling room-frame locations close to the relevant shock zone. These training and conflict maps are not correlated (r = −0.02, p = 0.67). The bar graphs quantify the comparison of room and arena positional information within 4.5 cm of the relevant shock zone normalized by the overall *I_pos_* and compared to other trial conditions. During training trials room-frame positional information is significantly higher near the training shock zone than during pretraining or conflict trials (F_2,8.9_ =8.7, p=0.008). During conflict trials room positional information is significantly higher near the relocated shock zone than during pretraining and training trials (F_2,9.8_ =9.0, p=0.006). Bar graphs represent mean ± SEM. * indicates different as shown by lines; * indicates significantly different from all other manipulations.

### Variable consistency of discharge coupling within the dentate network, specifically at sites of memory discrimination

Next, we investigated whether the sub-second coordination of DG spike trains changes across the trials requiring memory discrimination. The distributions of spike train correlations (τ) between pairs of pDGC place cells measured at the 133-ms time scale of the *I_pos_* estimates do not differ in the three trial types (H_2,381_ = 1.57, p = 0.46, Fig. S2.). Nor do they change systematically from one manipulation to another (Fig. 4A; H_2,381_ = 0.87, p = 0.65, Fig. S2). Standardized firing rates (z) were computed during 5-s intervals that identified passes through firing fields (Fenton et al., 2010; Fenton and Muller, 1998; Jackson and Redish, 2007). The distributions of standardized rates did not differ across the pretraining, training, and conflict sessions (F_2,9149_ = 0.31; p=0.73). Overdispersion of the standardized rates was not different in the pretraining sessions (var = 5.2) compared to values reported for area CA1 in rats. Overdispersion during pretraining was also indistinguishable from during training (var = 5.74; F_4189,3169_ = 1.10; p = 0.17), and conflict (var = 5.63; F_1791,3169_ = 1.08; p = 0.63) sessions (also when the unit of analysis was a single cell with at least 15 passes, F_2,178_ = 0.44; p=0.64). Similar to time-averaged measures of spatial firing (Fig. 2), single cell estimates of short time-scale spiking dynamics did not differ across the trials (Fenton et al., 2010; Fenton and Muller, 1998; Jackson and Redish, 2007).

**Fig. 4.**
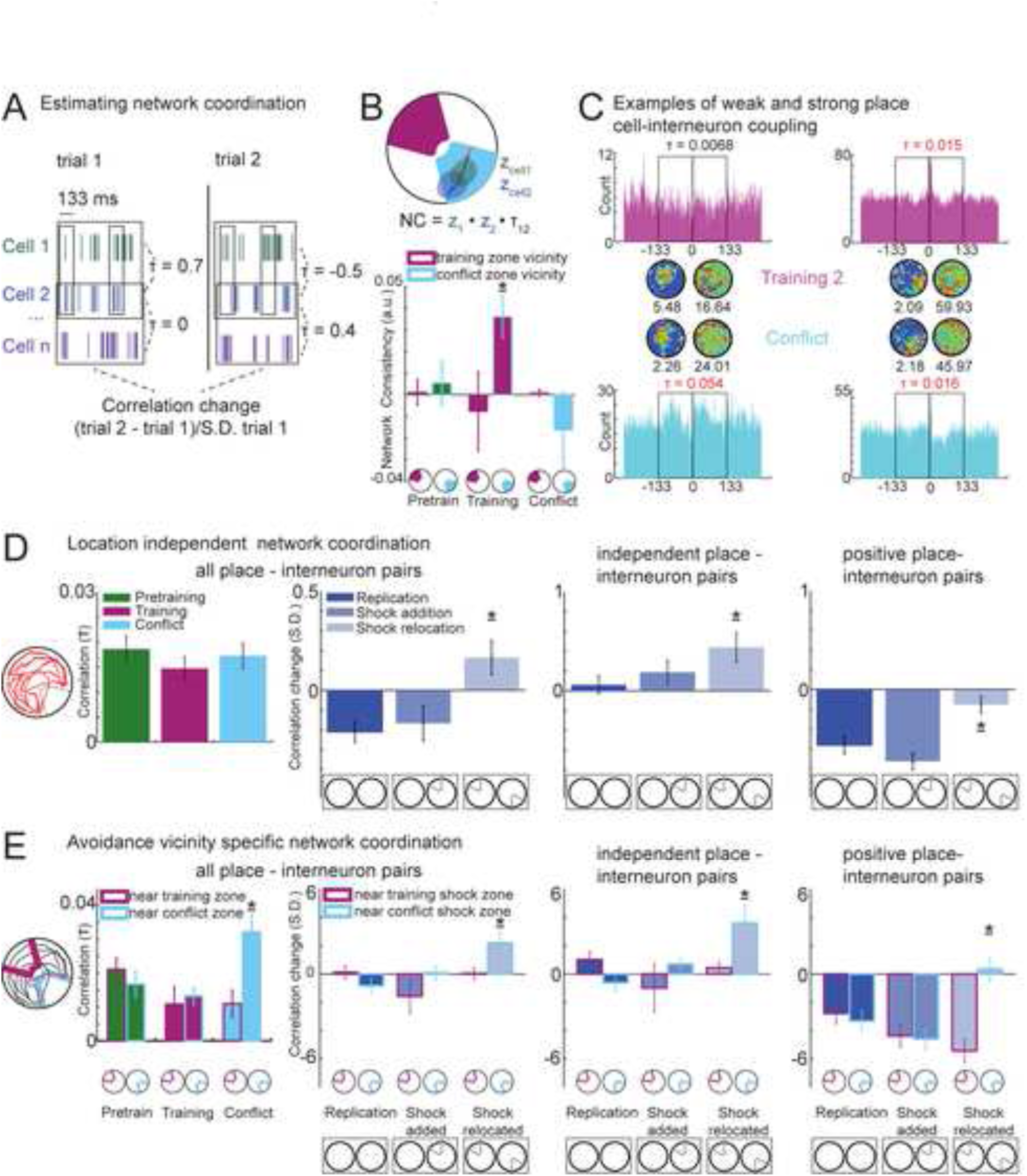
Sub-second coordination within the network of DG cells mirrors DG-dependent memory discrimination likelihood. a) Schematic of how pair-wise spike train correlation changes are estimated. Kendall’s correlation (τ) at 133 ms resolution is computed for all possible cell pair spike trains. The difference between the τ values for a pair of trials is normalized by the standard deviation of τ for that cell pair during the first trial. b) Top: cartoon of the momentary location-specific discharge of two neurons with overlapping place fields. Their momentary discharge relationship as the mouse passes across the firing fields (*z*_1_ · *z*_2_) is compared to their overall tendency to discharge together (*τ*_1,2_), and is measured as network consistency (*NC* = *z*_1_ · *z*_2_ · *τ*_1,2_). Bottom: Network consistency increases opposite the shock zone after training and decreases when the shock is relocated (shock zone: F_1,1038_=1.2, p=0.28, condition: F_2,1038_=1.5, p=0.22, shock zone x manipulation interaction F_2,1038_ = 3.2, p= 0.04). This indicates location-specific co-firing is less consistent at sites of memory discrimination. c) Firing rate maps of two example place cell-interneuron cell pairs and their cross-correlograms. (left) An uncorrelated cell pair during the training trial becomes correlated during the conflict trial, whereas (right) a positively correlated cell pair remains positively correlated. d1) No differences in correlations across conditions for place cell-interneuron pairs (H_2,435_ = 1.92, p=0.38). d2) During shock-relocation, place cell-interneuron pairs become more positively correlated compared to the replication and shock addition manipulations (F_2,435_ = 7.1, p<0.001). d3) The changes during shock-relocation, are due to the cell pairs that were initially independent becoming more positively correlated (H_2,236_ =8.1, p<0.017) while c4) the initially positively correlated pairs do not change. The initially-positive cell pairs decrease correlations during the replication and shock addition manipulations (H_2,280_ =24.3, p=10^−6^). Changes in correlated firing were specific to the vicinity of the shock zone. e1) Correlations are increased only when the animal is close to the relocated shock zone during the conflict trial (shock zone: F_1,806_ = 1.9, p = 0.15; condition: F_2,806_ = 2.6, p = 0.11; zone × condition: F_2,796_ = 6.7, p = 0.0013). The correlation increases demonstrated in panels d2-4 only occur when the animal is close to the relocated shock zone, when DG-dependent memory discrimination is most likely e2) all cell pairs: shock zone: F_1,746_=3.8, p=0.05; manipulation: F_2,746_= 6.3, p=0.002; shock zone × manipulation: F_2,731_= 5.5, p=0.0044; e3) independent cell pairs: shock zone: F_1,593_=1.3, p=0.3, manipulation: F_2,593_=4.4, p=0.01, shock zone x manipulation F_2593_ = 6.9, p= 0.0011; e4) positively-correlated cell pairs: shock zone: F_1,245_= 7.7, p=0.006; manipulation: F_2,245_= 2.9, p<0.06, shock zone x manipulation: F_2,245_ = 1 1.5, p<0.0001. The post-hoc tests on the data in panels e2-4 confirm that after shock relocation, the correlation changes near the relocated shock zone increase compared to the other zones and during the other manipulations. See also Figure S2. Bar graphs represent mean ± SEM. * indicates significantly different from all other manipulations.

We then considered the short time-scale spiking dynamics of place cell pairs. The momentary discharge of place cells with overlapping firing fields can characteristically covary positively so that when the mouse passes through the firing fields the two cells both fire more or less than predicted by their firing fields, or the cell pairs can covary negatively so when one fires excessively the other does not, or they can discharge independently so that the firing variations of one cell do not predict the other cell’s momentary firing (Fenton, 2015; Kelemen and Fenton, 2012). To evaluate the dynamic interactions amongst the network of spiking cells, we then asked whether the momentary firing rate variations of pairs of place cells as they crossed overlapping firing fields (*z_1_* and *z_2_*) were consistent with the overall short-time scale discharge coupling between the cell pair (*τ_1,2_*; see schematic in Fig. 4B top). Because *τ_1,2_* did not vary across conditions, we computed network consistency (*NC* = *z*_1_ · *z*_2_ · *τ*_1,2_), which is maximal when the momentary firing rates of two cells with overlapping firing fields covary similar to the overall spike train correlation of the cell pair; network consistency is minimal when the rate covariance is opposite to the spike train correlation. Although overall network consistency was positive and similar across the pretraining, training, and conflict sessions (F_2,2523_ = 13; p = 0.27), the requirement for memory discrimination predicts greater pattern distinctiveness and thus lower network consistency, particularly in the vicinity of the shock zone when memory discrimination is most required. During pretraining, before shock was ever experienced, network consistency was similar when the mouse was in the vicinity of the future shock zone and the opposite corresponding region of the arena (Fig. 4B bottom). During training trials, network consistency increased opposite the shock zone and network consistency in the same location decreased when the shock was relocated 180° in the conflict trial. While the increased network consistency opposite the initial shock zone was not predicted, the reduced network consistency in this location when shock is relocated is predicted by the greater demand for memory discrimination in the relocated shock zone vicinity, on the hypothesis that distinct network states manifest for distinct memories. Accordingly, momentary discharge amongst cell pairs will be inconsistent with the overall discharge coupling between network cell pairs (Fig. 4B bottom). This view can also explain increased network consistency opposite the initial shock zone if the mice consistently considered it safe and distant from shock. This pattern of change in network consistency can not simply be due to the mice spending more or less time in the shock zones because network consistency was unchanged across the session types in the vicinity of the initial location of shock, whether or not it was a neutral, avoided, of preferred region of the environment. We conclude that unlike session-averaged features of place cell discharge like the location of place fields, the sub-second discharge coordination within the network of pDGCs systematically varies across time and locations and these distinctive network states are associated with increased demand for memory discrimination and may be sensitive to rapid changes with learning experience (Bittner et al., 2015; Cheng and Frank, 2008).

### Increased place cell-inhibitory neuron coactivity at sites of memory discrimination

To explore how distinctive network states might transiently arise, we investigated whether the task manipulations alter discharge correlations between place cells and inhibitory neurons. Inhibitory networks may be important for memory discrimination (Buzsaki, 2010; Danielson et al., 2017; Jinde et al., 2013; Park et al., 2015), and network consistency, perhaps in part because high levels of inhibition promote pattern separation in mature granule cells (Marin-Burgin et al., 2012). The distributions of place cell – interneuron discharge correlations did not differ across trials (Fig 4C). In contrast, the correlated firing of the individual cell pairs differed across the three types of manipulation; the changes were higher across the shock-relocation manipulation (Fig. 4C top left) that requires memory discrimination and relies on intact DG function (Fig. 2). This increase in place cell – interneuron co-firing was observed generally, most place cells increased co-firing with at least one interneuron (83% of place cells for the shock-addition (z = 1.15, p = 0.2) and 92% for the shock-change (z = 2.3, p = 0.03) comparisons relative to 75% of place cells for the replication comparison. These findings also hold for 20 ms and 250 ms timescales (Fig. S2).

Because changes in correlated firing may not be homogeneous within a network of neurons (Harris, 2005; Okun et al., 2015; Olypher et al., 2006), the pairs were subclassified for further analysis. During pretraining, cell pairs were significantly positively (81 pairs, 53%), significantly negatively (15 pairs; 10%), or not (56 pairs, 37%) correlated according to statistical criteria. After shock-relocation, the correlations amongst the initially independent place cell-interneuron pairs increased (Fig. 4C top middle) whereas positively correlated pairs did not change, although they decreased with the replication and shock addition manipulations (Fig. 4C top right). Changes were not observed amongst the initially negatively correlated pairs (H_2,73_ =4.28, p = 0.12). These changes in correlated firing were specific to the vicinity of the shock zone, where memory discrimination is most required. The correlation increases are larger with the shock change when the mouse is close to the currently to-be-avoided zone than when it is close to the previous shock zone location (Fig 4C bottom). These location-specific differences were not detected for the other manipulations. In accord with these changes, during the conflict trial, we observed higher correlations near the current shock zone compared to its prior location (Fig 4C bottom). These differences between the locations were not observed when corresponding random samples of intervals were compared, nor were the location-specific differences observed in the pretraining or training trials. Together, these data indicate that coupling increases between the firing of DG place cells and inhibitory neurons, specifically when the animal needs to perform DG-dependent memory discrimination to avoid shock.

## Discussion

We evaluated hypotheses of how the dentate gyrus contributes to memory discrimination (Colgin et al., 2008; Leutgeb et al., 2007; Neunuebel and Knierim, 2014; Wills et al., 2005) by investigating the changes in pDGC place cell firing across different behavioral episodes that included a test demonstrated to require DG-dependent memory discrimination (Fig. 2A; Burghardt et al., 2012; Kheirbek et al., 2013). Contrary to expectations that time-averaged spatial discharge patterns (firing field arrangements) correspond to distinct spatial memories (Leutgeb et al., 2005a; Wills et al., 2005), we did not observe any form of firing field rearrangement (“remapping”) that was specific to the memory discrimination trial (Fig. 2F,G), even though spatial remapping of pDGC responses to spatial cue manipulations was greater than of Ammon’s horn (Fig. 1B; Leutgeb et al., 2007). We also detected that firing rates in and out of place fields varied across cue manipulations (Fig. 1C) and changed task contingencies (Figs. 2G). Thus, the responses to cue manipulations replicate prior reports. In contrast, place fields did not remap with the memory discrimination test. This is in opposition to the idea that distinctive spatial discharge patterns underlie spatial memory discrimination, but resembles how the place cell network responds to displaced objects; by changing which object-coding cells co-fire with which place cells, none of which remap (Muller and Kubie, 1987; Rivard et al., 2004). Instead of memory discrimination triggering remapping, we found that DG-dependent memory discrimination is signaled by network inconsistency: weaker correspondence between momentary co-firing amongst cell pairs with overlapping spatial tuning and overall sub-second discharge correlations (Fig. 4). The memory task-related variations in network consistency and the corresponding modulation of excitatory - and inhibitory cell coupling suggest that memory task-related network representations in DG place cell firing are distinguished by sub-second interactions between excitatory discharge signaling spatial information and local control of that discharge by interneurons (Buzsaki, 2010; Danielson et al., 2017). Because the set of weak pair-wise correlations within a network estimates the overall network state (Schneidman et al., 2006) the present observations demonstrate that the network of cells in the DG adopts distinctive states defined by dynamic and coordinated excitatory-excitatory and excitatory-inhibitory interactions to subserve memory discrimination. Like the requirement for memory discrimination, the coordinated discharge interactions were dynamic, and could be best estimated by analysis of momentary activity rather than by session-averaged measurements. The findings (Fig. 4C-E) indicate that the sub-second co-firing of dentate principal cells and interneurons, transiently increases during moments of difficult memory discrimination, pointing to greater interneuron-driven control of dentate network function during difficult memory discriminations (Fig. S2D). These distinctive memory-associated patterns of network discharge presumably feedforward to CA3 to implement memory discrimination. This network view is based on recordings that do not discriminate between granule and mossy cells in the DG, which may have distinctive extracellular discharge properties (Danielson et al., 2017; GoodSmith et al., 2017; Neunuebel and Knierim, 2012; Senzai and Buzsaki, 2017). Mossy cells reside in the hilus of the DG and receive focal input from local granule cells (Amaral, 1978; Scharfman et al., 1990). In addition to exciting granule cells directly, mossy cells excite DG inhibitory neurons broadly along the dorso-ventral axis that then provide strong global inhibition onto granule cells and may consequently establish the lateral-inhibition component of the functional architecture for a competitive network (Amari, 1977), specialized for discriminating input patterns and network states (Scharfman, 2016; Sloviter and Lomo, 2012). Correspondingly, ablating mossy cells causes hyperexcitability in granule cells and impaired pattern separation (Jinde et al., 2013) and according to a recent computational model loss of mossy cell excitatory drive onto granule cells does not affect pattern separation, while loss of the mossy cell driven inhibition of granule cells impairs pattern separation (Danielson et al., 2017). While the present physiological findings cannot distinguish between differential roles of granule cells and mossy cells in memory discrimination, nonetheless a crucial contribution of these DG cells was demonstrated using optogenetic silencing (Fig. 2B; Kheirbek et al., 2013). We were unable to functionally discriminate between the pDGC recordings by dividing them into distinct classes on the basis of features that may discriminate granule cells and mossy cells, including firing field number, firing rate, spike width and preferred theta phase (GoodSmith et al., 2017; Senzai and Buzsaki, 2017). Nonetheless, the findings point to a network perspective that focuses on the interactions amongst granule, mossy, and inhibitory cells to emphasize the network state of the DG rather than the isolated contribution of single cell classes. Future work will do well to define the integrated roles of these classes by determining if specific cell classes make particular contributions that merit distinctive functional classification. Future work should also evaluate the distinctiveness of inputs in comparison to the pDGC output to critically evaluate the notions of pattern separation that have been assigned to these cells. In summary, substantial evidence from behavioral tests have established an important role of the DG in memory discrimination, which we find is mediated by changes in the network state of sub-second interactions amongst excitatory and inhibitory cells within the DG, rather than by the spatial tuning of principal cells measured across the time scales of minutes, much longer than the time scale of memory discrimination and decision.

## Author contributions

MvD and AF designed the experiments. MvD collected and analyzed the data. MvD and AF wrote the paper.

## Acknowledgements

Supported by NIH grant R01AG043688. We want to acknowledge Younghun Lim and Zejia Angel Yu for help with experiments. We are grateful to Gyorgy Buzsaki, René Hen, and Helen Scharfman for discussions, guidance and comments on the manuscript.

## Supplemental experimental procedures

### Subjects

Nine C57BL/6 and sixteen Cre-POMC-Halorhodopsin male adult mice were used (8 Cre+ and 8 Cre-). All procedures and care adhered to protocols approved by New York University Animal Welfare Committee, which follow NIH guidelines. Cre-POMC-Halorhodopsin mice result from the following cross: B6.FVB-Tg(Pomc-cre) 1L owl/ J (RRID : IM SR_JAX:010714) x B6;129S-Gt(ROSA)26Sortm39(CAG-hop/EYFP)Hze/J (RRID:IMSR_JAX:014539).

### Surgery

Mice were anesthetized with isoflurane (3% for induction 1.5-1.75% maintenance) for surgical procedures. Ketoprofen post-surgical analgesic was administered for three days after surgery. A minimum of 2 weeks passed before experimental procedures began.

### Optic Fiber implantation

Bilateral optic fibers (0.125 diameter optic fibers (NA 0.37) in 1.25mm zirconia ferrules) were surgically implanted at ML: ±1.25, AP: −1.8, DV −1.55 using C&B Metabond to attach the ferrules to the skull.

### Electrode implantation

Custom microdrives with 4 independently movable tetrodes (17μm tungsten wires, gold plated to ~120 kΩ impedance at 1 kHz) were surgically implanted to center bregma coordinates ML:1.25, AP: −1.5, DV: −1.2. Sterile vaseline covered the tetrode shafts and C&B Metabond and Grip Cement (Densply) was used to attach the microdrive to the skull and bone screw. A bone screw was placed at ML:-3, AP +0.5 relative to bregma, and served as electrical ground. Mice recovered from surgery for at least 1 week before lowering the tetrodes.

### Recording set up and single unit analysis

A unity-gain preamplifier configured as a source-follower was connected to the mouse and the signals were conducted to a commercial recording system (Axona Ltd., St. Albans, U.K.). The signals were filtered (300 Hz - 7 kHz), amplified 2000 - 8000 times, and digitized at 48 kHz for recording presumptive action potentials using the dacqUSB system (Axona Ltd., St. Albans, U.K.). Local-field potentials were filtered (0.1 - 200 Hz), amplified (100-1000 times) and digitized (2 kHz). The position of the mouse was determined 30 times a second by tracking two LEDs attached to the preamplifier using video tracking (Tracker, Bio-Signal Group Corp., Acton, MA). The spatial resolution at which the location-specific discharge of cells was analyzed is 1.5 cm/pixel. Action potential waveforms were analyzed offline using custom Wclust software (A.A. Fenton). Single unit discrimination quality was objectively assessed using IsoI metrics (Neymotin et al., 2011); average *IsoI_background_* = 6.52±0.15 bits; average *IsoI_nearest neighbour_* = 7.58±0.23 bits. Firing rate maps were computed by dividing the number of action potentials that a cell discharged in a location by the amount of time the mouse was detected in that location. Unitary waveforms during the first recording trial of the experimental sequence were assigned to putative cell classes according to the following features: waveform width, firing rate, firing rate map information content, coherence, and % active pixels.

### Numerical Analyses and statistics

Analyses were performed using custom-written Matlab code (2015B, Mathworks). One-way and two-way repeated measures ANOVAs were used as appropriate. When data were not normally distributed or the group variances were unequal, one of the following non-parametric tests was used, as appropriate: two-way ANOVA on ranks, Kruskal-Wallis or Welch’s one-way ANOVA. Fisher’s LSD were performed for post-hoc pair-wise comparisons.

### Place cell responses to environmental manipulations

Seven mice explored three versions of an environment. 1) Standard: a transparent 40-cm diameter circular enclosure with three cue cards on the wall that was placed within a 50 x 50 cm square box also with a cue card; 2) Rotated: the same circular enclosure in the box with all cue cards rotated 90°; 3) Removed: the square box by itself after removing the circular enclosure. Three comparison conditions were evaluated by comparing place cell properties across a pair of recordings conditions: 1) Replication compared standard-standard, rotated-rotated (if available), and removed-removed (if available) conditions; 2) Rotation compared standard-rotated conditions; and 3) Removal compared standard-removed, and rotated-removed conditions.

### Active place avoidance

A commercial active place avoidance apparatus and software was used (Bio-Signal Group, Corp., Acton, MA). Mice were trained on the active place avoidance task in a black-curtained room with distinctive visual cues on the curtains. For optogenetic silencing, a 593 nm laser was attached to the implanted optic fibers. When the laser was on the peak power in the brain was 15 mW. On day 1, mice explored the rotating disk during a 30-minute pretraining trial with no shock. On day 2 mice learned to avoid a computer-controlled 0.2 mA, 60 Hz, 500 ms foot shock in a 60° zone during two 30-minute training trials (2 hours apart). The shock zone was stationary with respect to the room (Cimadevilla et al., 2001). On the third trial of day 2 the mice were trained in a conflict trial with the shock zone relocated 180°. Mice were euthanized and perfused 105 minutes after the start of the conflict trial.

Eight transgenic POMC-Halorhodopsin that express the light sensitive chloride channel halorhodopsin selectively in DG granule cells and eight littermate controls received the active place avoidance training as described above.

Six mice implanted with tetrodes also performed the active place avoidance task, but with five sessions that took place during one day with 90 min rest periods after each session. In two 20 min pretraining sessions, the mice explored the rotating disk while the action potential discharge of DG units were recorded. Next in a 20 min training session mice learned to avoid a 60° shock zone. Immediately after the training session there was a 20 min recording session with no shock to avoid electrical artifacts, during which the mice continued to avoid the shock zone. Immediately after the recording session there was a 5 min reminder with shock on in case the mice had begun to extinguish the avoidance. The second training trial began after a 90 min rest in the home cage. The mice had a 10-min training trial with the shock on, after which there was a 20 min recording session with shock off. After another 90 min rest period the mice received conflict training for 20 min with the shock zone relocated 180°. Immediately after the conflict training there was a 20 min recording session with shock off.

### Firing rate map stability

Spatial remapping was estimated by the firing rate map stability, measured as Fischer’s z-transformation of the Pearson correlation. The correlation was computed for the corresponding pixels from the firing rate maps from the two recordings. The measure was taken to be the maximum correlation after allowing for one map to move with respect to the other a maximum of 2 pixels in the x and y dimensions. This ensured that small changes due to experimental error were minimized. The active place avoidance trials produce spatial behavior and single unit discharge in two spatial frames. Firing rate maps from both spatial frames were analyzed and the higher correlation was taken as the estimate for map stability. If there were multiple trial correlations for a given manipulation (e.g. standard 1 - rotated, standard 2 - rotated) the average correlation was used to estimate the value for each cell.

Firing rate remapping was estimated as the difference between a cell’s average rates in two trials divided by the maximum of the two rates. Place field rate remapping was estimated similarly; the field firing rate in each session was defined by a “field mask”, pixels that comprised the largest place field in the first trial of the day. We also calculated the corresponding change in the out-of-field rate by estimating the average rate in all the “non-mask” pixels that did not include the field mask. Thus separate measures of spatial remapping, rate remapping, firing field remapping, and out-of-field remapping were evaluated.

### Positional information

Positional information (*I_pos_*) was calculated for each place cell at 133 ms temporal resolution (Kelemen and Fenton, 2010; Olypher et al., 2003):

I_pos_=|*p(n\x)log2(p(n\x)/(p(n))*|, where p(n|x) is the probability of observing *n* action potentials in 133 ms at location *x*, and *p*(*n*) is the probability of observing *n* action potentials in 133 ms independent of location. *I_pos_* during the arena rotation was frame specific because it was separately computed for positions in the room and arena spatial frames.

Ensemble *ΔI_pos_* was calculated by independently summing the room frame and the arena frame *I_pos_* time series of cells in an ensemble and subtracting the two frame-specific *I_pos_* sums at each 133 ms time step, such that a positive value indicated a momentary preference for room frame information and a negative value indicated an arena frame preference (Kelemen and Fenton, 2010). During each trial, the total amounts of time room- and arena-preferring ensemble discharge were observed was estimated as the total time Δ*_pos_* was positive and negative, respectively. Local minima of the mouse’s distance to the nearest edge of the shock zone were calculated and selected if they were closer to the shock zone than the animal’s average position. The *I_pos_* values during the ~1 s prior to the minima estimated frame-specific discharge during an active avoidance. This was calculated for each mouse as the average of the *Ipos* values preceding each minimum. This average was computed for the vicinity of each shock zone location on each trial (even when there was no shock zone as in pretraining trials). Summary maps of the proportion of time that discharge was room-preferring in each pixel were calculated for each mouse and averaged over all the animals. To estimate the frame-preference of spatial discharge in the vicinity of the shock zone, the room and arena *I_pos_* values were calculated separately and averaged for those times during which the mice were within 3 pixels (4.5 cm) of the shock zones or in the shock zone itself.

### Cell pair spike train discharge correlations

Kendall’s correlation was calculated for all possible place cell pairs in an ensemble and for all place cell-interneuron pairs (interneuron-interneuron pair numbers per animal were not sufficient for analysis). The correlation was computed at the 20 ms, 133 ms, and 250 ms time scales; the data from the 133 ms time scale are reported in the main text and the conclusions were the same for the other time scales. How a cell pair’s discharge correlation changed across two recordings was estimated by subtracting the correlation in one trial from the correlation of that cell pair in the other trial. The difference was divided by the standard deviation of the correlation to avoid non-linear estimates of correlation changes due to correlations being near zero. To estimate the standard deviation of a cell-pair’s discharge, the discharge correlation was calculated six times after dividing the earlier trial into six periods. The standard deviation was based on the six correlation estimates. Kendall’s correlation was also calculated for the time intervals that the mouse was within 3 (4.5 cm) pixels of the shock zones but not in the zones themselves.

### Overdispersion

Overdispersion measures the reliability of a place cell’s spatial discharge (Fenton and Muller, 1998). It assumes that the session-averaged firing rate map is an accurate prediction of the momentary discharge of a cell as the subject moves across a firing field. The expected firing during a pass through a firing field is determined on the basis of the Poisson assumption, by summing the product of the rate (*r_x_*) and the duration (*t_x_*) in each location (*x*) of the N locations that define the pass: *expected rate* = 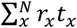. Then for each pass we can compute the standardized rate as:

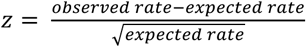. If the observed firing is much more or less than expected from the session-averaged firing rate map, the variance of the distribution of *z’s* will exceed two standard deviation units, and then the cell’s discharge is overdispersed (Jackson and Redish, 2007; Olypher et al., 2002).

Standardized rates were computed as in (Fenton et al., 2010) by computing *z* for each 5-s episode, during which *exp* was greater than the average firing of the cell. Only passes with locations (*x*) that had been sampled more than 0.67 s were considered.

### Network Consistency

Network consistency measures the reliability of a pair of place cells’ spatial co-firing. It assumes that the cell pair’s session-wide correlated discharge is an accurate prediction of the momentary co-firing of the cell pair as the subject moves across the overlapping firing fields of the two cells. The standardized firing of each cell of the pair along the pass (*z_1_ andz_2_*) is computed as described for overdispersion, above. Kendall’s correlation (t_1,2_) was used to estimate the overall tendency of the cell pair to co-fire during a short time interval (e.g. 133 ms). Network Consistency on a single pass was computed as:

*NC* = *z*_1_ · *z*_2_ · *τ*_1,2_. The product of the two standardized rates during an individual pass was used as an independent estimate of how the discharge of the cell pair covaries. If the momentary discharge across passes covaries like the overall correlation for the cell pair then *NC* is maximized and if the momentary discharge covaries opposite to the overall correlation, then *NC* is minimized. As in computing overdispersion, the standardized rate on the pass for each cell estimates the momentary variation of firing from the session-averaged expectation:

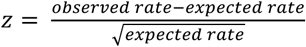. It the expected rate was smaller than the session-averaged rate of that cell then the pass was excluded to ensure that only passes through a place field were evaluated (Fenton et al., 2010). To estimate zone specific network consistency, we added an additional constraint by only calculating the standardized rates on passes through firing fields in the vicinity of each potential shock zone (including the surrounding 6 cm). Only passes through the shock zone vicinity that were longer than 3 s were considered. The distribution of *NC* was analyzed and compared between recording conditions.

### Histology

Mice were anesthetized with a sodium pentobarbital solution (100mg/kg, i.p.) and perfused with saline followed by 4% paraformaldehyde in 0.1 M phosphate buffer (transgenic and control mice) or 10% formalin (electrophysiology mice). The brains were removed and post-fixed overnight. To identify the electrode tracks, mice heads with the brain partially exposed were put in formalin solution overnight, after which the brains were removed and post-fixed for another day. After post-fixing, the brains were cryoprotected in 30% sucrose 0.1 M phosphate buffer solution. Fifty μm sections through the hippocampus were stained by cresyl violet to locate the electrode tracks. The sections were 35 μm for the transgenic and control mice and the tissue was stained with c-Fos Anti-Rabbit primary antibody (1:8000; from Millipore) and 594 nm Goat Anti-Rabbit Alexa Fluor (1:500) secondary for fluorescent microscopy. The sections were mounted with Vectashield mounting medium with DAPI (Vector Labs). To estimate neuronal activity during the conflict trial c-Fos+ cells were counted in six slices per animal for 4 Cre+ and 4 Cre-control mice.

**Figure S1.**
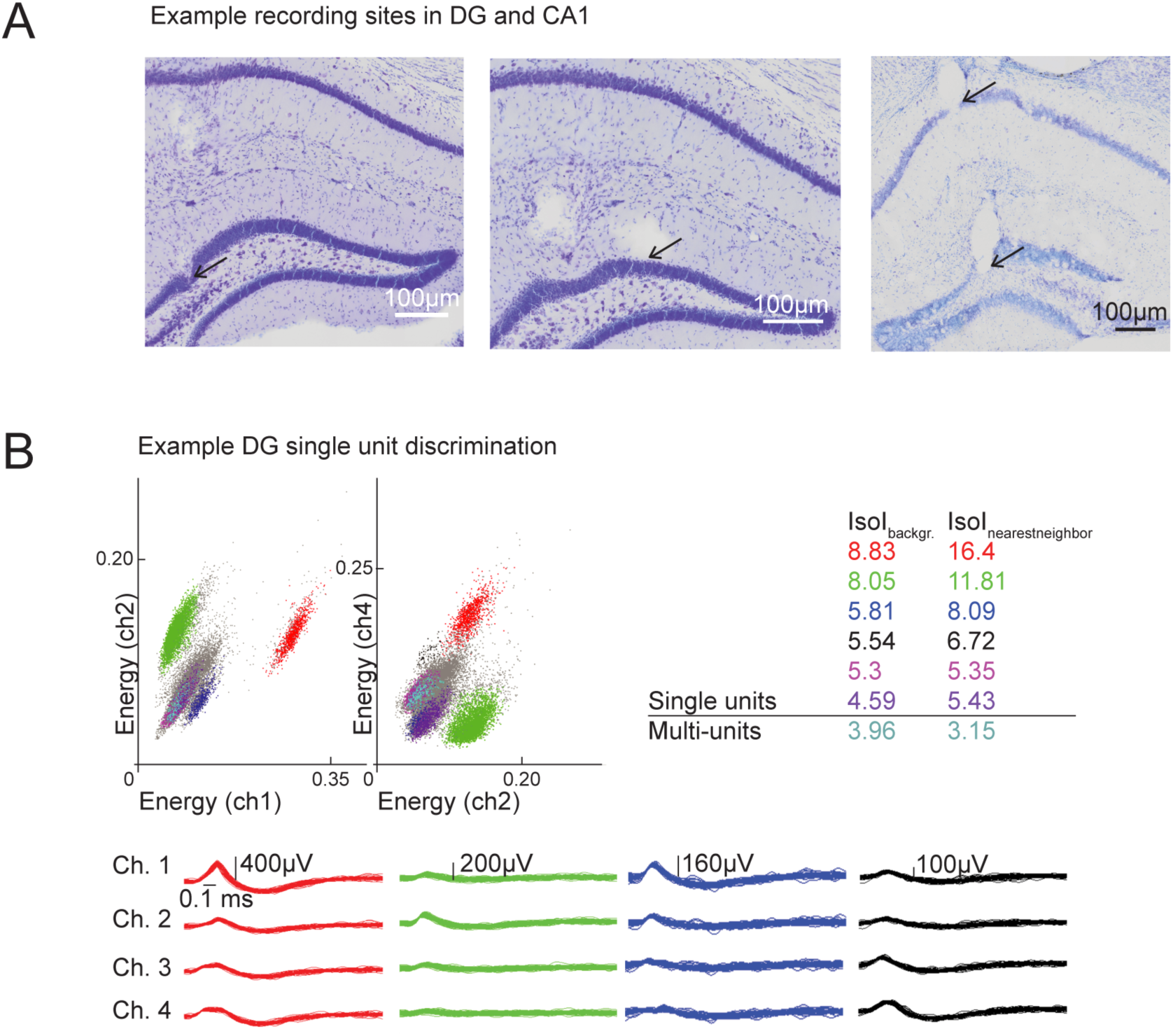
Example recording sites and single units. Related to Figure 1. a) Examples of recording sitesin DG and CA1. B) Example illustrating single unit discrimination of action potential wave shapes. Related to Figure 1.

**Figure S2.**
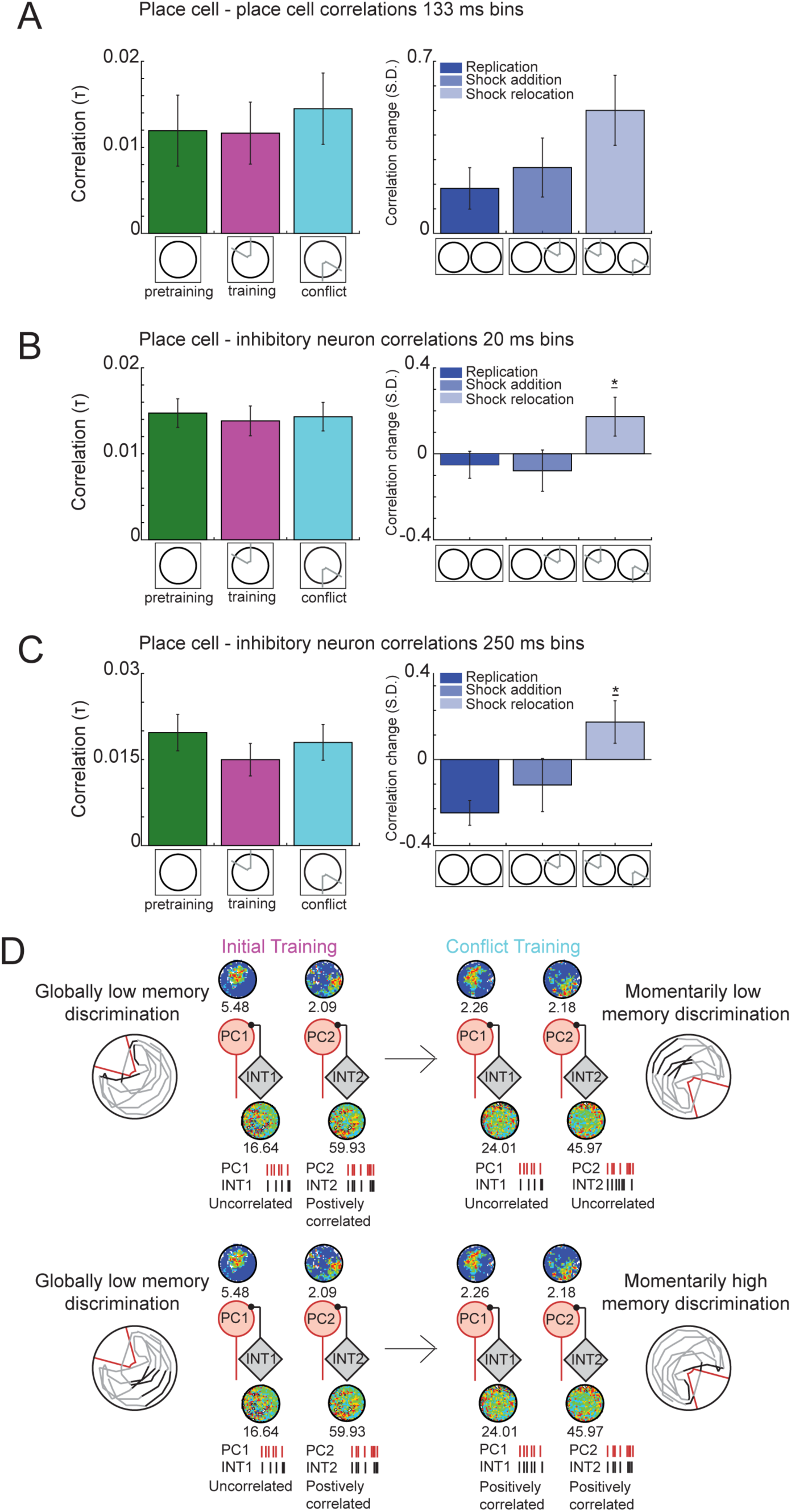
Cell-pair spike train discharge correlations. Related to Figure 4. a) Place cell-place cell discharge correlations computed at 133 ms resolution. Left: No differences between the trial types (H_2,381_ = 1.57, p = 0.46). Right: No differences in correlation changes between the manipulations (H_2,381_ = 0.87, p = 0.65). b) Place cell-interneuron discharge correlations at a 20 ms resolution. Left: No differences between the trial types (H_2,435_ = 0.29, p = 0.86). Right: During DG dependent memory discrimination in the shock relocated manipulation, place cell-interneuron pairs become more positively correlated compared to the replication and shock addition manipulations (H_2,435_ = 6.17 p=0.046). c) Place cell-interneuron discharge correlations at 250 ms resolution. Left: No differences between the trial types (H_2,435_ = 1.59, p = 0.45). Right: During DG dependent memory discrimination in the shock relocated manipulation, place cell-interneuron pairs become more positively correlated compared to the replication and shock addition manipulations (H_2,435_ = 17.18 p=10^−4^). d) Schematic model of DG network activity during DG-dependent memory discrimination. Although the conjoint discharge of place cell – interneuron cell pairs is characteristic for each cell pair (uncorrelated, positively or negatively correlated), place cell - interneuron cell pairs increase co-firing during moments of difficult memory discrimination, pointing to greater control of location-specific ensemble discharge during difficult memory discriminations. Related to Figure 4.

**Table S1.** Place cell characteristics for first session in the standard environment. Related to Figure 1.

| Place cell characteristics | DG |  | CA |  |
| --- | --- | --- | --- | --- |
|  | Average | S.E.M. | Average | S.E.M. |
| **0.3 peak width (μs)** | 204 | $\pm 7.29$ | 193 | $\pm 8.54$ |
| **firing rate (AP/s)** | 1.04 | $\pm 0.18$ | 1.00 | $\pm 0.16$ |
| **% ISI <10ms** | 0.22 | $\pm 0.02$ | 0.28 | $\pm 0.04$ |
| **map coherence (z)** | 0.53 | $\pm 0.03$ | 0.55 | $\pm 0.05$ |
| **info content (bits/s)** | 1.41 | $\pm 0.12$ | 1.30 | $\pm 0.18$ |
| **mean ISI (ms)** | 5.99 | $\pm 0.59$ | 3.93 | $\pm 0.60$ |
| **# place fields** | 1.42 | $\pm 0.11$ | 1.26 | $\pm 0.13$ |

**Table S2.** Place cell characteristics for three active place avoidance task conditions. Related to Figure 2.

| Place cell characteristics | Pretraining |  | Training |  | Conflict |  |
| --- | --- | --- | --- | --- | --- | --- |
|  | Average | S.E.M. | Average | S.E.M. | Average | S.E.M |
| **0.3 peak width (μs)** | 10.34 | $\pm 0.34$ | 10.04 | $\pm 0.33$ | 10.07 | $\pm 0.38$ |
| **firing rate (AP/s)** | 0.99 | $\pm 0.17$ | 0.87 | $\pm 0.13$ | 0.82 | $\pm 0.16$ |
| **% ISI <10ms** | 0.23 | $\pm 0.02$ | 0.23 | $\pm 0.02$ | 0.21 | $\pm 0.02$ |
| **mean ISI (ms)** | 5.94 | $\pm 0.97$ | 6.05 | $\pm 0.91$ | 5.31 | $\pm 0.45$ |
| **info content room (bits/s)** | 1.6 | $\pm 0.12$ | 1.45 | $\pm 0.12$ | 1.75 | $\pm 0.18$ |
| **# fields room** | 2.36 | $\pm 0.21$ | 1.99 | $\pm 0.18$ | 1.81 | $\pm 0.21$ |
| **p active pixels arena** | 0.46 | $\pm 0.03$ | 0.40 | $\pm 0.03$ | 0.37 | $\pm 0.03$ |
| **map coherence arena (z)** | 0.33 | $\pm 0.02$ | 0.30 | $\pm 0.03$ | 0.25 | $\pm 0.02$ |
| **info content arena (bits/s)** | 1.41 | $\pm 0.11$ | 1.43 | $\pm 0.12$ | 1.70 | $\pm 0.18$ |
| **# fields arena** | 2.94 | $\pm 0.27$ | 2.80 | $\pm 0.28$ | 2.68 | $\pm 0.30$ |

